# Primary human chondrocytes respond to compression with plasma membrane receptors and by microtubule activation: a phosphoproteomic study

**DOI:** 10.1101/672352

**Authors:** Donald L. Zignego, Jonathan K. Hilmer, Brian Bothner, William J. Schell, Ronald K. June

## Abstract

Chondrocytes are responsible for maintaining the cartilage that helps joints like the knee and hip bear load and move smoothly. These cells typically respond to physiological compression with pathways consistent with matrix synthesis, and chondrocyte mechanotransduction is essential for tissue and joint homeostasis. In osteoarthritis (OA), chondrocyte mechanotransduction appears to be dysregulated, yet many pathways and mechanisms of osteoarthritic chondrocyte mechanotransduction remain poorly understood. The objective of this study is to document the phosphoproteomic responses of primary osteoarthritic chondrocytes to physiological sinusoidal compression. Here we show that OA chondrocytes respond to physiological compression by first activating proteins consistent with cytoskeletal remodeling and decreased transcription, and then later activating proteins for transcription. These results show that several microtubule-related proteins respond to compression, as well as proteins related to calcium signaling, which has previously been extensively shown in chondrocytes. Our results demonstrate that compression is a relevant physiological stimulus for osteoarthritic chondrocytes. We anticipate these data to be a starting point for more sophisticated analysis of both normal and osteoarthritic chondrocyte mechanotransduction. For example, finding differences in compression-induced phosphoproteins between normal and OA cells may lead to druggable targets to restore homeostasis to diseased joints.

## Introduction

Osteoarthritis (OA) is the most common joint disorder worldwide [1–7], and is characterized by the breakdown of the articular cartilage covering the joint surfaces. Articular cartilage is composed of a dense extra cellular matrix (ECM), a less-dense pericellular matrix (PCM), and highly specialized cells, chondrocytes [8]. At these joint surfaces (*e.g*. the hip and knee), the chondrocytes are subjected to repetitive mechanical loading, which can reach magnitudes as high as 10 times an individual’s body weight [9]. These loads alter the chondrocyte environment. In OA, the processes by which chondrocytes sense and respond to their mechanical environment, termed mechanotransduction [10], are disrupted.

Chondrocytes [10–13], and other mammalian cells [14–17] can transduce mechanical inputs into biological signals, but the link between these two processes remains unclear [18]. Because most of the druggable genome resides in phosphoprotein-mediated interactions [19], understanding how the chondrocyte phosphoproteome changes in response to compression is important for developing new treatments for joint disease. The objective of this study is to characterize changes in protein phosphorylation after dynamic compression of primary osteoarthritic chondrocytes.

Chondrocytes are the sole cell type in articular cartilage and play a critical role in maintaining cartilage homeostasis through anabolic and catabolic processes. Mechanical stimulation drives this delicate balance [20, 21]. The role of healthy chondrocytes is primarily anabolic in nature. This anabolism includes protecting, maintaining, and repairing cartilage by synthesizing collagen (mostly type II), and proteoglycans [22] through the secretion of cytokines, growth factors, and protease inhibitors [20]. However, in diseased cartilage (*e.g*. OA), catabolism dominates, and usually involves breakdown of ECM and PCM through secretion of proteases (*e.g*. matrix metalloproteinase (MMPs)). Dynamic loading, such as walking, promotes anabolic responses in diseased chondrocytes, whereas static loading inhibits matrix anabolism (*e.g*. by upregulation of catabolic enzymes such as MMP-13) [23–26].

Previous studies of signaling in chondrocytes discovered pathways involving proliferation [27], cell differentiation and dedifferentiation [28], matrix catabolism (via MMPs and ADAM/ADAM-TS gene expression) [29], and programmed cell death [30]. These studies provide a detailed understanding of various processes for mechanically-induced signaling in chondrocytes. Studies of chondrocyte mechanotransduction show that chondrocytes respond to applied loading by remodeling their cytoskeleton [31, 32]. Previous phosphoproteomic analysis of primary human chondrocytes provided insight into the pathophysiology of degradative diseases in cartilage [22]. However, to our knowledge, no studies have used phosphoproteomics as a tool for elucidating signaling mechanisms of chondrocyte mechanotransduction.

In this paper, we move toward understanding the role of protein phosphorylation in chondrocyte mechanotransduction. To do this, we identify phosphorylated proteins in primary human OA chondrocytes that are dynamically compressed to physiological levels. The data show a phosphoproteomic signature consistent with mechanotransduction beginning at the cell membrane and proceeds through the cytoskeleton to the nucleus. This spatiotemporal sequence of pathways can inform studies aiming to harness chondrocyte mechanotransduction to repair and rebuild cartilage.

## Results

The objective of this study is to characterize the phosphoproteomic response of primary human OA chondrocytes stimulated by physiological dynamic compression. We analyze the untargeted phosphoproteomic profiles to minimize bias by the exclusion of important, but unexpected, data (*i.e*. if only a single signaling pathway were selected *a priori*). Primary chondrocytes from OA joint replacement tissues were harvested, encapsulated in physiologically stiff agarose, and subjected to physiological compression tests for 15 (DL15) or 30 (DL30) minutes prior to phosphoproteomic analysis (Figure 1). We also analyzed uncompressed control samples (UC). To our knowledge, the present study is the first application of phosphoproteomics to analyze mechanotransduction in primary OA chondrocytes. These data demonstrate that applied dynamic compression stimulates primary osteoarthric chondrocytes to alter their phosphoproteomic profiles in a 3D culture system.

**Figure 1.**
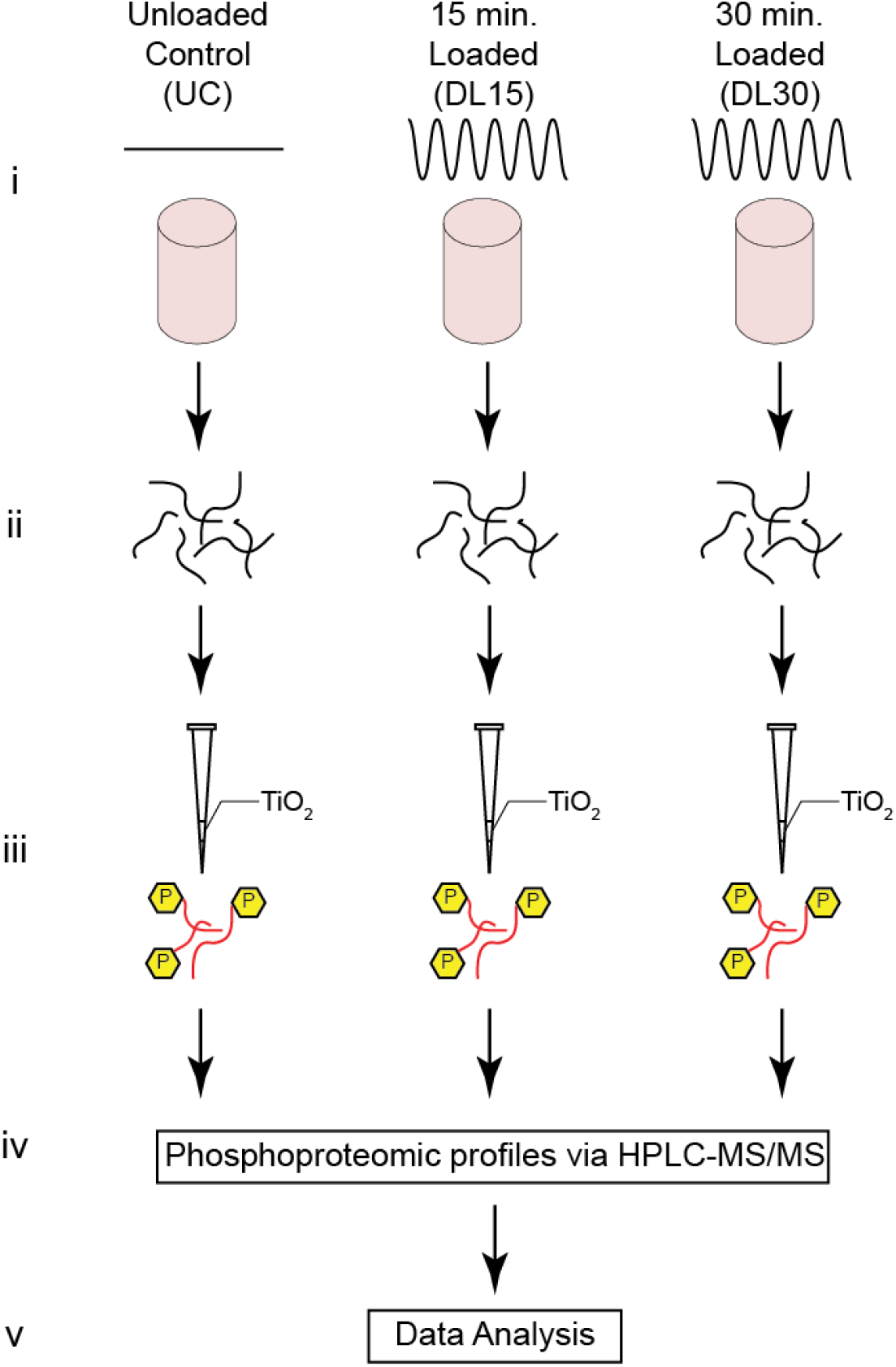
Experimental Design. (A) Schematic for both untargeted experimental methods. (i) Primary human OA chondrocytes are encapsulated in physiologically stiff agarose (4.5% agarose, stiffness ~35 kPa), cultured for 72 hours, and then dynamically compressed in tissue culture for 0, 15, or 30 minutes (Control, DL15, or DL30) at 1.1 Hz. (ii) Proteins are extracted by flash freezing the samples, pulverizing, and lysing the cells followed by overnight enzymatic digestion. (iii) Samples are enriched for phosphopeptides using TiO_2_ enrichment, (iv) phosphoproteomic profiles identified via HPLC-MS/MS, and (v) the untargeted data analyzed.

### Proteomic Analysis and Characterization

LCMS data for each of the individual samples (n=5 per group) from each experimental group (UC, DL15, and DL30) were processed individually through the TOPPAS pipeline [33]. After false discovery rate (FDR) filtering with a 5% threshold and using the decoy protein hits and combining all replicates from each loading group, we identified 2858, 2246, and 2570 phosphoproteins including isomers for UC, DL15, and DL30, respectively (Supplementary Table 6). We further refined these data by focusing only on phosphoproteins detected in more than half of the samples for each loading group. The final dataset consisted of 767, 359, and 623 phosphoproteins detected for UC, DL15, and DL30, respectively (Figure 2). To analyze the effects of dynamic compression on chondrocyte metabolism, we made individual comparisons between unloaded control samples and samples loaded for either 15 or 30 minutes of dynamic compression, discussed below. We first report differences in phosphoprotein expression before examining differences in pathways represented by various groups of proteins.

**Figure 2.**
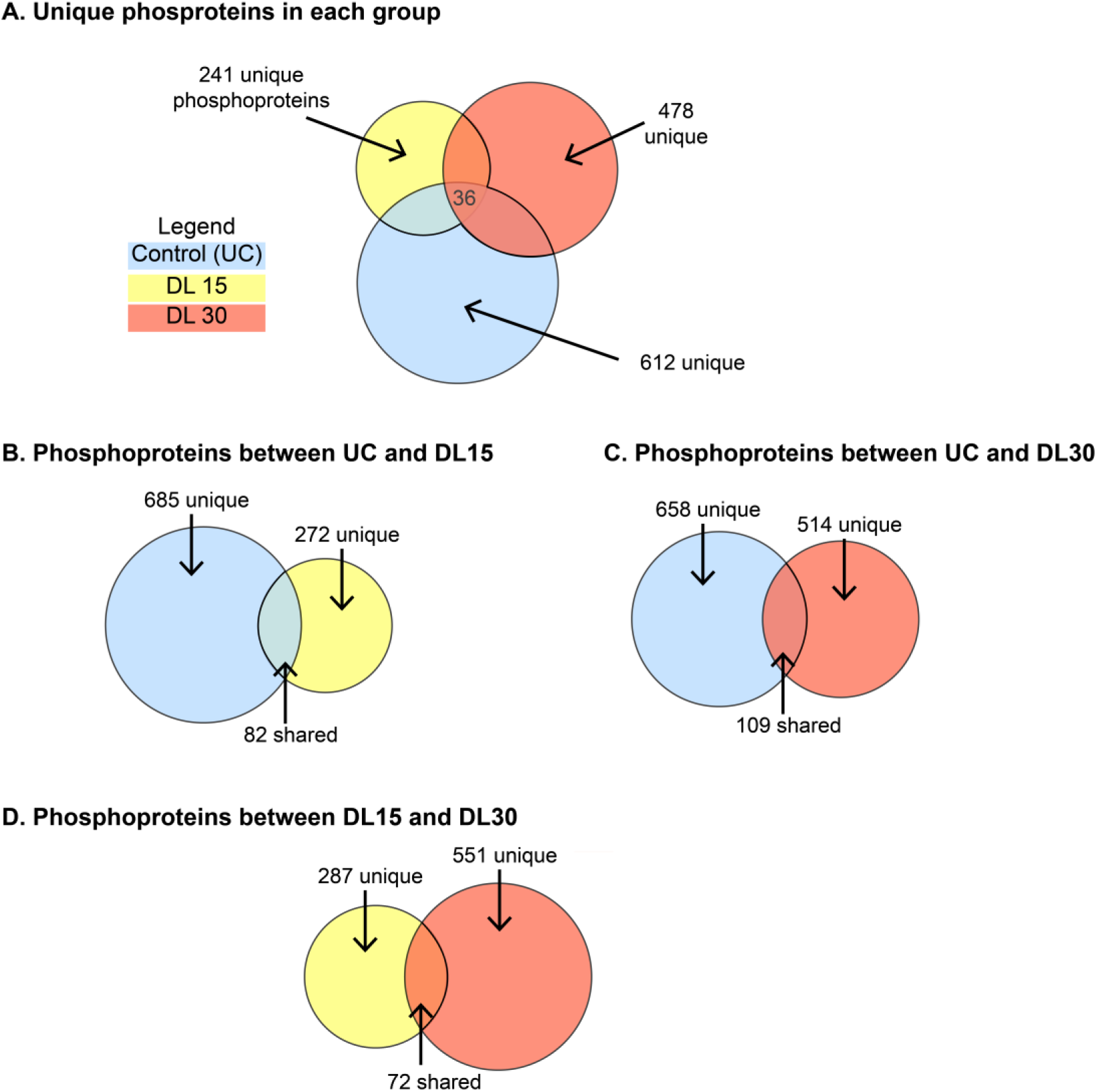
Dynamic compression alters phosphoprotein in 3D primary OA chondrocytes. To determine signaling pathways activated by dynamic compression, 3D-encapsulated OA chondrocytes were encapsulated in agarose and subjected to 0 (control, UC), 15 minutes (DL15), or 30 minutes (DL30) of compression before analysis by LC-MS that examined phosphoprotein isomers based on detected peptides. Figure quantifies detected proteins using Venn Diagrams with circle areas proportional to the number of proteins. Unique refers to the number of proteins within the indicated area. (A) Unique patterns of phosphoproteins were observed in each group. (B) There were 277 unique phosphoproteins detected after 15 minutes of compression in comparison to unloaded controls. (C) There were 514 phosphoproteins detected after 30 minutes of compression in comparison to controls. (D) There were both unique and shared phosphoproteins detected between 15 and 30 minutes of compression. Proteins shown in File S5.

### Differential phosphoprotein expression after 0, 15, or 30 minutes of compression

By comparing samples stimulated by dynamic compression to the controls, we identified proteins phosphorylated in response to mechanotransduction. This group helped us to establish the breadth of compression as an environmental stimulus. Comparing all groups (e.g. UC, DL15, and DL30, Figure 2A), we find 612 phosphoproteins are unique to UC, 241 unique to DL15, 478 unique to DL30, and 36 common between all three of them. The first comparison between unloaded controls (UC) and DL15 samples reveals 685 phosphoproteins unique to UC, 277 unique to DL15, and 82 phosphoproteins common between the two samples (Figure 2B).

Of the 82 shared phosphoproteins, 2 are significantly (p<0.05) up-regulated by mechanical loading (E3 ubiquitin-protein ligase UBR4 and Coiled-coil domain-containing protein 87) and 1 is downregulated (Protein PRR14L). Levels of the remaining 79 proteins are not significantly different. For the second comparison, we compare UC against DL30 samples (Figure 2C). 658 phosphoproteins are unique to UC, 514 unique to DL30, and 109 common between both samples. Of the 109, 4 are up-regulated and 2 down-regulated (all p<0.05). The remaining 103 that were shared, were not significantly different. For the third comparison (DL15 vs. DL30, Figure 2D), 287 phosphoproteins are unique to DL15, 551 unique to DL30, and were 72 common between them. Of the 72 common between DL15 and DL30, one phosphoprotein is down-regulated with increased loading in the DL30 group and the remaining 71 are not significantly different.

We identify three unique clusters of phosphoproteins that were co-regulated as a result of dynamic compression using unsupervised clustering (Figure 3, Table 1). These clusters describe the timecourse of protein phosphorylation stimulated by compressive loading. Cluster 1 includes phosphoproteins up-regulated with only 15 minutes of dynamic compression. Cluster 2 contains phosphoproteins down-regulated as a result of dynamic compression (either 15 or 30 minutes), and cluster 3 identified phosphoproteins up-regulated only after 30 minutes of dynamic compression. Phosphorylated proteins from cluster 1 include microtubule cross-linking factor 1 (MTCL1), unconventional myosin-Va (MYO5A), ankyrin-2 (ANK2), and obscurin (OBSCN). Cluster 2 phosphorylated proteins include cofilin-1 (COF1), microtubule-actin crosslinking-factor 1 (MACF1), E3 ubiquitin-protein ligase (UBR4), and calreticulin (CALR). Phosphorylated proteins from Cluster 3 include vimentin (VIME), unconventional myosin-IXb (MYO9B), Titin (TITIN), Protein AF-9 (AF9), mediator of RNA polymerase II transcription subunit 13 (MD13L), and Vam6/Vps39-like protein (VPS39). Taken together, these data show a broad response to compression. This includes cytoskeletal regulation, regulation of intracellular calcium, organelle trafficking, and transcription through RNA polymerase II.

**Figure 3.**
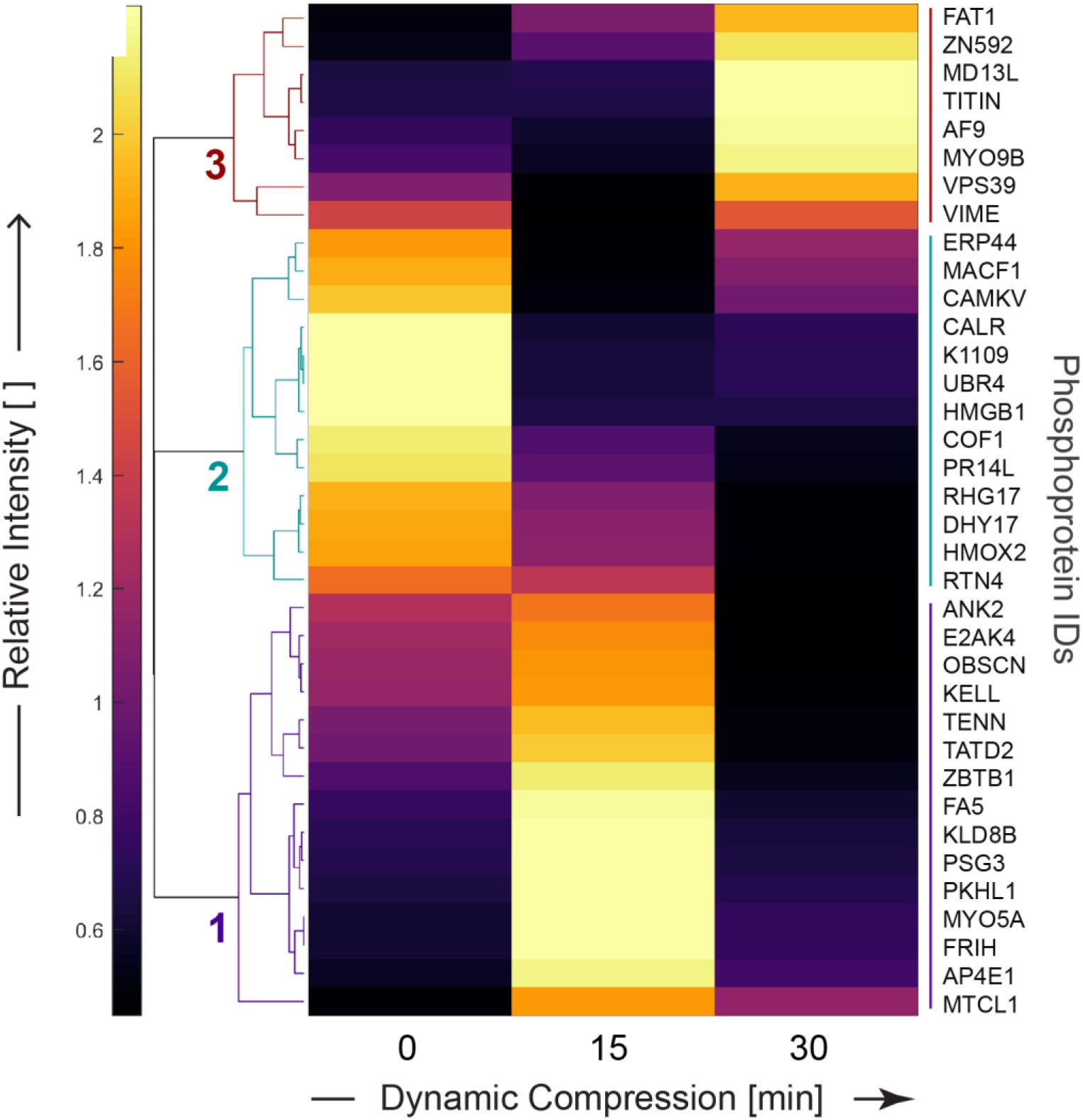
Applied compression resulted in distinct untargeted phosphoproteomic profiles for primary OA chondrocytes. Clustering of phosphoprotein intensities common to all groups (e.g. intersection from Fig. 2A) found three groups of mechanically regulated protein phosphorylation. Patterns of expression displayed by heatmap following unsupervised, hierarchical clustering on differentially phosphorylated proteins common to the dynamic compression (15 and 30 minutes) and unloaded control groups. Heatmap labels are protein identifiers, with complete names found in Supplementary Table 5.

**Figure 4.**
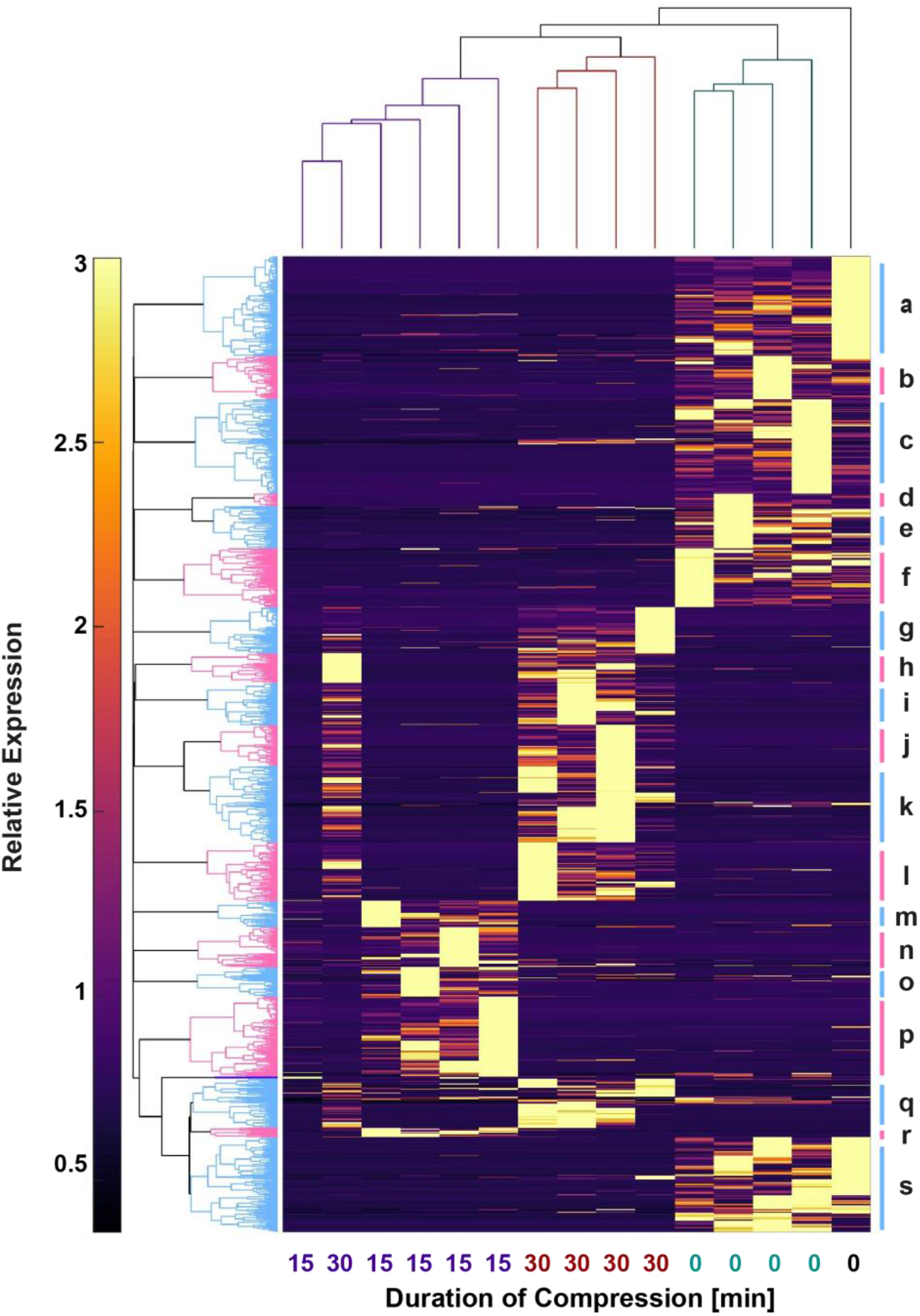
Clustering indicates strong effects of mechanical loading on global phosphoprotein expression. Unsupervised clustering of all phosphoproteins detected in all samples conducted in MATLAB. Vertical dendrogram (top) indicates that the uncompressed controls (UC) were the most distinct samples, while there was similarity between the samples compressed 15 and 30 minutes. Individual clusters of phosphoproteins labeled alphabetically beginning on the top right. Pathways associated with these clusters shown in Supplementary File Pathway_Enrichment_Results.zip. Key cluster results shown in Table 3.

**Table 1.**
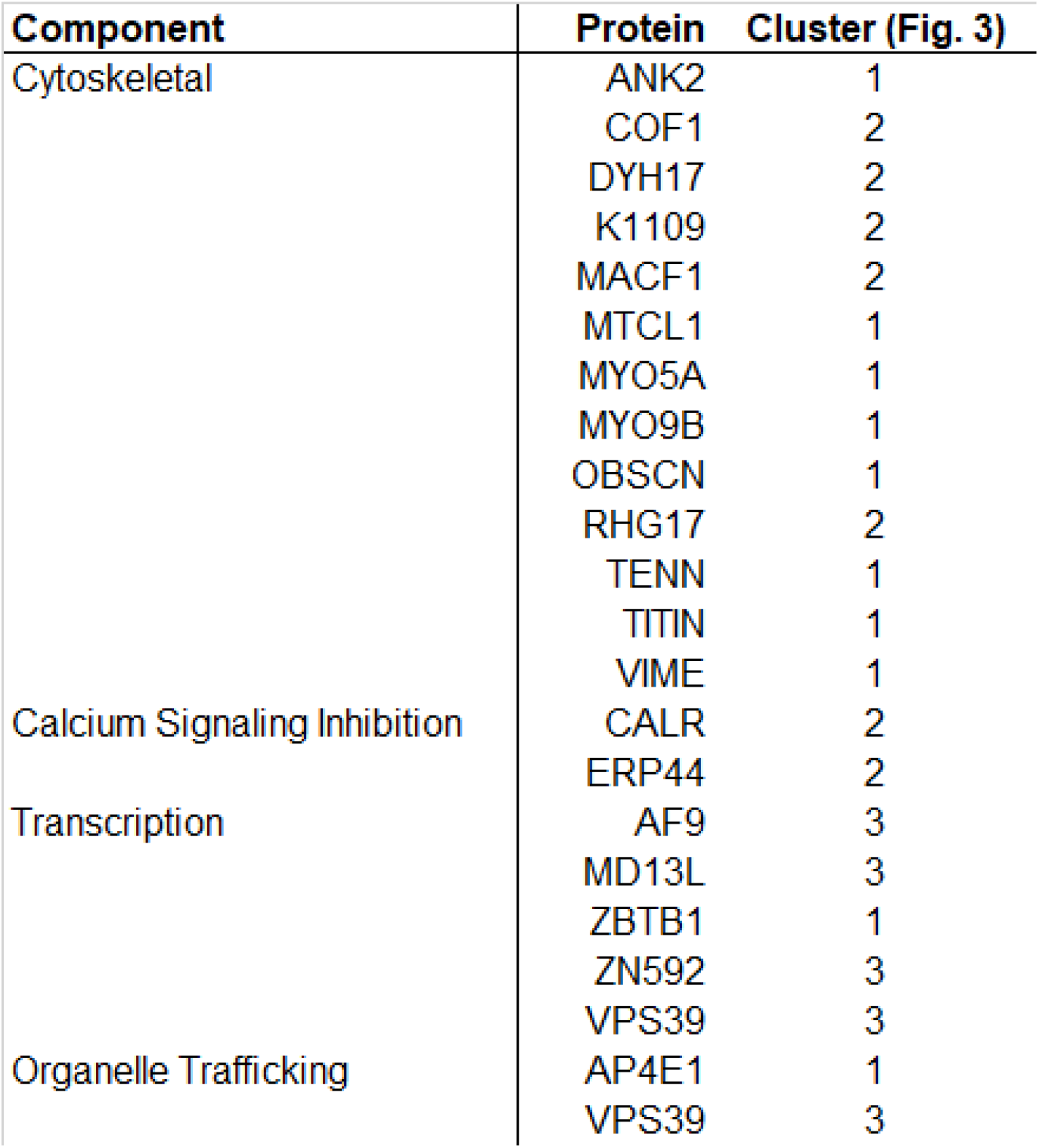
Key proteins common to all groups suggest mechanotransduction involves cytoskeletal, calcium regulation, organelle trafficking, and transcriptional elements. Table summarizes key proteins detected in all groups from clusters in Figure 3.

These data show that dynamic compression substantially alters expression of phosphorylated proteins, and that there are differences between cells stimulated by 15 and 30 minutes of compression. To further understand chondrocyte mechanotransduction, we analyzed pathways represented by these phosphoproteins.

### Pathways stimulated by 0, 15, or 30 minutes of compression

To examine how compression regulated the chondrocyte phenotype, we use pathway enrichment analysis. The phosphoproteins identified after FDR-correction from each of the comparisons (UC vs. DL15, UC vs. DL30, and DL15 vs. DL30) identify key signaling pathways using enrichment analysis (Table 2, Supplemental Tables 2-4) [34]. This analysis compares the relative enrichment of phosphoproteins between specific signaling pathways and the specific phosphoproteins identified in each group.

**Table 2.** Pathways affected by physiological compression of OA chondrocytes. Over-representation analysis performed using IMPALA based on proteins detected in clusters in Figure 4. Pathways defined as significant when p-values were smaller than the FDR-corrected significance level of 0.05. PID = pathway interaction database. INOH = Integrating Network Objects with Hierarchies. Wiki = wikipathways. Reactome = reactome.

| Observation | Pathway (cluster from Fig 4) | Database | Pathway Proteins |  | Proteins (Uniprot IDs) | P-value |
| --- | --- | --- | --- | --- | --- | --- |
|  |  |  | # Detected | Total |  |  |
| Increased by 15 minutes compression | Regulation of CDC42 activity (p) | PID | 4 | 40 | GIT1; DOC10; BCAR3; DOCK9 | 7.48E-05 |
|  | Urokinase-type plasminogen activator (uPA) and uPAR-mediated signaling (o) | PID | 3 | 33 | DOCK1; FIBG; LRP1 | 1.67E-04 |
| Increased by 30 minutes compression | Arginine Proline metabolism (h) | INOH | 3 | 54 | KCRS; ARG1; AOC3 | 2.03E-04 |
|  | FTO Obesity Variant Mechanism (j) | Wiki. | 2 | 8 | PRD16; IRX3 | 3.01E-04 |
| Inhibited by compression (higher in control) | GP1b-IX-V activation signalling (a) | Reactome | 3 | 10 | VWF; GP1BA; FLNA | 6.31E-05 |
|  | Peptide hormone biosynthesis (a) | Reactome | 3 | 14 | INHBE; NEC1; COLI | 1.87E-04 |
|  | Hedgehog (c) | INOH | 5 | 72 | PRS6B; GLI2; PRS4; PSMD2; KIF3A | 1.97E-04 |
|  | Senescence and Autophagy (s) | Wiki. | 6 | 105 | IBP5; CDN1B; LAMP1; SQSTM; CO1A1; IBP3 | 2.41E-04 |
|  | FOXM1 transcription factor network (c) | PID | 4 | 42 | CCNA2; XRCC1; NFAC3; CENPF | 2.63E-04 |
|  | IL-6 signaling (f) | INOH | 2 | 6 | IL6RB; JAK2 | 3.54E-04 |
|  | Collagen chain trimerization (s) | Reactome | 4 | 44 | CO1A1; CO1A1; CO1A2; COOA1 | 4.81E-04 |
|  | RNA Polymerase III Transcription (d) | Reactome | 2 | 42 | TF3B; BDP1 | 9.65E-04 |
|  | Intestinal lipid absorption (d) | Reactome | 1 | 1 | NPCL1 | 1.11E-03 |

When comparing UC against DL15, we observe 201 signaling pathways (p<0.05) unique to UC samples. These pathways included collagen formation and biosynthesis (p<0.01). 92 significant pathways are enriched from phosphoproteins unique to DL15 samples, including Acetyl-CoA biosynthesis (p=0.0157) and Hedgehog signaling (p=0.0386). Phosphoproteins common to both UC and DL15 show 61 significant signaling pathways, including RhoA activity (p<0.01), and Rho GTPase signaling (p<0.01) that have previously been observed in chondrocyte mechanotransduction [35]. These data indicate that 15 minutes of loading induces phosphoproteomic changes at the cell periphery.

For the UC vs. DL30 comparison, we identify 215 signaling pathways (p<0.05) unique to UC, 119 unique to DL30, and 71 shared by both UC and DL30. The pathways for UC include collagen formation and biosynthesis (p<0.01), and extracellular matrix organization (p=0.01). Pathways for DL30 samples included MAPK signaling (p<0.01), Rho GTPase signaling (p<0.01), hyaluronan biosynthesis (p=0.0252), and glucose-6-phosphate dehydrogenase deficiency (p=0.0473). Overlapping pathways between UC and DL30 included ECM proteoglycan synthesis (p=0.0374), and Erk2 activation (p=0.0489). These data indicate that 30 minutes of compression activate a pattern of signals consistent with matrix synthesis (e.g. hyaluronan) and nuclear signaling (e.g. MAPK).

To examine how compression changed over time, we compare DL15 to DL30. We find 80 signaling pathways unique to DL15, 100 unique to DL30, and 90 shared by both DL15 and DL30 (all p<0.05). Specific pathways to DL15 samples include Hedgehog (p<0.01), and calcium signaling (p=0.0453), consistent with prior studies [36, 37]. Specific pathways to DL30 samples include MAPK signaling (p<0.01), fatty acid activation (p<0.01), and Rho GTPase signaling (p<0.01). Common pathways between both DL15 and DL30 samples included RhoA activity (p=0.0168) and nucleotide sugars metabolism (p=0.0342). These data emphasize the role that RhoA plays in responding to applied compression of chondrocytes. Taken together, they support the concept of mechanotransduction beginning at the membrane (*e.g*. calcium signaling) and moving toward the nucleus (*e.g*. MAPK) with potential feedback for membrane remodeling by fatty acid activation through Acetyl-CoA metabolism.

### Protein domains affected by compression

To assess sequence similarities between proteins affected by dynamic compression of chondrocytes, protein domain analysis was performed using DAVID [38, 39] using FDR-correction. There were several structural domains common to the uncompressed controls (Table 3). These include several helicase domains indicating that dynamic compression deactivates helicases. Furthermore, domain analysis of proteins detected after either 15 or 30 minutes of compression indicates that Dynein, myosin, ATPase, and Calmodulin-binding domains are activated by compression.

**Table 3.** Protein domains absent after compression. Proteins unique to uncompressed controls were analyzed for structural domains using DAVID. Helicase domains are enriched in these proteins, indicating that dynamic compression deactivates helicases, which are required for de novo transcription.

| Protein Domain | Accession # | Database | P-value |
| --- | --- | --- | --- |
| Armadillo-like fold. | IPR016024 | Interpro | 2.63E-02 |
| ATP-binding domain, helicase superfamily 1-2 | IPR014001 | Interpro | 6.77E-03 |
| C-terminal domain, helicase superfamily 1-2 | IPR001650 | Interpro | 5.44E-03 |
| DEAD-like helicases superfamily | SM00487 | SMART | 4.76E-02 |
| Helicase conserved C-terminal domain | PF00271 | Pfam | 2.74E-02 |
| Superfamilies 1 and 2 helicase ATP-binding type-1 domain profile | PS51192 | Prosite | 1.15E-02 |
| Superfamilies 1 and 2 helicase C-terminal domain profile | PS51194 | Prosite | 1.15E-02 |

**Table 4.** Protein domains detected after compression. Proteins present after either 15- or 30-minutes of compression were analyzed for structural domains using DAVID.

| Protein Domain | Accession # | Database | P-value |
| --- | --- | --- | --- |
| AAA - ATPases | SM00382 | SMART | 1.65E-03 |
| AAA domain (dynein-related subfamily) | PF07728 | PFAM | 2.46E-04 |
| AAA+ superfamily ATPase domain | IPR003593 | INTERPRO | 2.79E-04 |
| ATPase, dynein-related, AAA domain | IPR011704 | INTERPRO | 2.26E-04 |
| ATPase; molecular moto | SM00242 | SMART | 2.34E-02 |
| Calmodulin-binding motif. | SM00015 | SMART | 5.19E-03 |
| Dynein heavy chain | IPR026983 | INTERPRO | 2.76E-05 |
| Dynein heavy chain and region D6 of dynein motor | PF12777 | PFAM | 3.06E-05 |
| Dynein heavy chain region D6 P-loop domain | IPR004273 | INTERPRO | 2.76E-05 |
| Dynein heavy chain, coiled coil stalk | IPR024743 | INTERPRO | 2.76E-05 |
| Dynein heavy chain, domain-1 | IPR013594 | INTERPRO | 8.06E-03 |
| Dynein heavy chain, domain-2 | IPR013602 | INTERPRO | 2.76E-05 |
| Dynein heavy chain, N-terminal region 1 | PF08385 | PFAM | 8.39E-03 |
| Dynein heavy chain, N-terminal region 2 | PF08393 | PFAM | 3.06E-05 |
| Dynein heavy chain, P-loop containing D4 domain | IPR024317 | INTERPRO | 4.12E-04 |
| Dynein motor, microtubule-binding stalk | PF12777 | PFAM | 3.06E-05 |
| isoleucine-glutamine calmodulin-binding motif | PF00612 | PFAM | 1.26E-02 |
| isoleucine-glutamine motif | PS50096 | PROSITE | 1.46E-02 |
| isoleucine-glutamine motif, EF-hand binding site | IPR000048 | INTERPRO | 1.16E-02 |
| Myosin head (motor domain) | PF00063 | PFAM | 1.09E-02 |
| Myosin head, motor domain | IPR001609 | INTERPRO | 9.73E-03 |
| Myosin heavy chain signature | PR00193 | PRINTS | 7.91E-03 |
| Myosin motor domain | PS51456 | PROSITE | 1.25E-02 |
| P-loop containing dynein motor region D4 | PF12780 | PFAM | 4.47E-04 |
| P-loop containing nucleoside triphosphate hydrolases | SSF52540 | SUPFAM | 2.22E-06 |
| P-loop containing nucleotide triphosphate hydrolases | 3.40.50.300 | GENE3D | 9.23E-03 |
| P-loop containing nucleotide triphosphate hydrolases | IPR027417 | INTERPRO | 1.09E-07 |

## Discussion

We used phosphoproteomics to identify pathways used by primary chondrocytes to respond to compression. During OA, cartilage deterioration proceeds through catabolic processes within the joint. Matrix degradation softens and weakens cartilage, and further joint loading through physiological processes (*e.g*. walking) promotes additional joint damage. Because chondrocytes and all cells respond to mechanical stimuli [40], these loads also provide an opportunity for therapeutic direction of mechanotransduction to restore cartilage and joint health. These data show that even chondrocytes from joints with advanced grade IV osteoarthritis can respond to applied compression with a broad phosphoproteomic response.

Here we found phosphorylation changes consistent with calcium signaling, cytoskeletal remodeling, and modulation of transcription and in response to applied compression of primary OA chondrocytes. We analyzed the short-term (<30 min) phosphoproteomic changes of these chondrocytes by comparing unloaded control samples (0 min. of loading), to samples subjected to either 15 (DL15) or 30 (DL30) minutes of dynamic compression. These data complement previous metabolomic findings [41], and expand basic knowledge of chondrocyte mechanotransduction. Cytoskeletal changes mainly appeared after 15 minutes of loading whereas transcriptional regulation followed these changes after 30 minutes of loading. Importantly, these cytoskeletal changes likely begin prior to the 15 minute timepoint observed in this study [35].

Calreticulin was de-phosphorylated in response to applied compression. Calcium signaling is a well-established mechanosensitive response [42–44] and there are interactions between the actin cytoskeleton and calcium signaling [44–46].

The cytoskeleton is a dynamic structure that provides crucial physical linkages within the cytosol and across the plasma membrane. We observed phosphorylation of ankyrin-2 and actin after 15 minutes of dynamic compression. We further observed dephosphorylation of the E3 ubiquitin ligase UBR4 which promotes actin remodeling [47]. After 30 minutes, we observed phosphorylation of vimentin, a perinuclear intermediate filament [48]. Finally, regulation of CDC42 activity was an enriched pathway after 15 minutes of compression. These data suggest “outside-in” mechanotransduction, wherein an externally applied load is first (*e.g*. after 15 minutes in this study) perceived at the plasma membrane and is then transduced through intermediate filaments which act as secondary transducers of mechanical loads.

Ankyrin not only affects chondrocyte cytoskeletal dynamics, but is important for maintaining cartilage homeostasis through CD44 [49]. This provides a potential mechanism whereby CD44 binds extracelluar hyaluronan for ECM deformation to be transmitted intracellularly. The actin cytoskeleton is a key component of cell shape, which has been linked with matrix synthesis supporting the chondrocyte phenotype [50, 51], and prior studies found active remodeling of chondrocyte actin in response loading [12, 51, 52]. Disruption of vimentin networks in chondrocytes leads to decreased matrix synthesis and cell stiffness [53], and OA chondrocytes exhibit altered vimentin networks [48].

The cytoskeleton contains microtubules. However, little is known about the role of microtubules in chondrocyte mechanotransduction. Here, we found that compression decreased phosphorylation of microtubule-actin crosslinking factor 1 (MACF1) and dynein heavy chain 17 (DYH17), while compression increased expression of microtubule crosslinking factor 1 (MTCL1.) DYH17 produces force at the minus end of microtubules and MACF1 crosslinks microtubules to actin [54]. MTCL1 helps organize microtubules during polarization.

Furthermore, several dynein heavy chain and ATPase domains were present in proteins detected after both 15 and 30 minutes of compression. Additionally, both adaptor protein complex 4ε1 (AP4E1) and VPS39 were increased after compression. These proteins are involved in organelle traffic and sorting, potentially indicating compression-dependence in these processes. Taken together, these results suggest that compression induces microtubule reorganization. However, the effects of this reorganization (*e.g*. increased cellular stiffness, organelle trafficking, or other effects) remain unclear.

Cytoskeletal signaling regulates transcriptional activity [55]. Consistent with our interpretation that early (e.g. before 15 minutes) compression-induced cytoskeletal changes activate later transcriptional responses, after 30 minutes we observed phosphorylation of AF-9. AF-9 is a component of the RNA Pol II complex that accelerates elongation of nascent mRNA precursors [56]. Also after 30 minutes of compression, Cam6/Cps39-like protein, a regulator of TGF-beta activity through activation of SMAD2-dependent transcription, was phosphorylated. Furthermore, we also observed compression-induced phosphorylation of zinc finger proteins (ZN592 and ZBTB1). ZBTB1 is a transcriptional repressor [57], which was elevated after 15, but not 30 minutes of compression. ZNF592 contains the KRAB (Kruppel-associated box) domain which also represses transcription [58, 59]. Taken together, these data suggest that compression induces a specific transcriptional profile that includes repression of target genes for ZN592 and ZBTB1, consistent with several prior studies [23, 60].

To our knowledge, this is the first study to perform proteome-wide phosphorylation analysis following applied compression to chondrocytes encapsulated in a physiologically stiff 3D microenvironment [61, 62]. There are important limitations to this study. First, this study focused on phosphoproteins, and therefore the expression of many unphosphorylated proteins is likely compression-dependent. This phosphoproteomic response is consistent with compression-dependent changes in transcription, and future transcriptional profiling may identify the specific genes induced by applied compression. Finally, compressive-sensitive phosphoproteomic expression may be affected by variables not considered in this study such as pericellular matrix, cell age, and other types of mechanical load such as shear.

In summary, this study demonstrates the power of using phosphoproteomics as a tool for understanding the chondrocyte response to short-duration (<30 min), dynamic compression. By expanding upon prior metabolomic analysis of primary human OA chondrocytes in response to dynamic compression [41], we identified 514 phosphoproteins unique to dynamically stimulated samples. To our knowledge, this was the first study to successfully identify phosphoproteomic profiles for OA human chondrocytes in response to mechanical loading. This work suggests the potential to use mechanical stimulation (*i.e*. short-duration, low-impact exercise) as a tool to promote cartilage repair in OA clinical populations.

## Materials and Methods

### Chondrocyte Culture and Encapsulation

Primary human chondrocytes were harvested, isolated, and encapsulated using previously optimized methods [61–63]. Chondrocytes were harvested from the femoral heads of n=5 Grade IV OA patients undergoing total hip joint replacement surgery (mean age: 63 years (range: 54-80), and mean mass: 80.4 kg (range: 56.9-99 kg)). To harvest chondrocytes, cartilage shavings were digested in Type IV collagenase (2 mg/mL for 12-14 hrs. at 37°C), and cultured in DMEM with 10% fetal bovine serum and 1% antibiotics (10,000 I.U. /mL penicillin and 10000 μg/mL streptomycin) in 5% atmospheric CO_2_. After 1 passage, cells were encapsulated at a concentration of ~500,000 cells/gel, and equilibrated in tissue culture conditions for 72 hours.

### Mechanical Stimulation

Mechanical stimulation was performed using established techniques [41, 63]. For each donor (n=5, female), cell-seeded agarose gels were randomly assigned to one of three loading groups: unloaded controls (*i.e*. 0 minutes of loading), 15, or 30 minutes of dynamic, cyclic compression (n = 5 biological replicates for each loading group). Homogenous deformations [62] were applied to cell-seeded gels using previously optimized methods [41, 63] with a custom-built bioreactor emulating physiological loading conditions (3.1-6.9% compressive strain, calculated from initial gel height and a frequency = 1.1 Hz [64]).

### Protein Preparation and Extraction

To extract proteins after compression, gels were flash frozen in liquid nitrogen, pulverized, and stored at −80°C prior to cell lysis [41, 63]. Cells were lysed by sonication and vortexing in RIPA buffer (50mM Tris-HCL (pH 8.0), 50mM NaCl, 1% NP-40, 0.5% sodium deoxycholate, and 0.1% SDS [65]). Samples were then centrifuged at 21,000 x g at 4°C for 10 minutes. Proteins were precipitated from the supernatant using ice-cold acetone overnight at −20°C. Samples were centrifuged the next day at 21,000 x g at 4°C for 10 min, and the acetone supernatant was discarded. The purified protein pellet was then re-suspended in 0.5M triethylammonium bicarbonate (TEAB).

### Proteolysis, TiO_2_ Phosphopeptide Enrichment, and Graphite Cleanup

To identify phosphoproteins, we digested the proteins as follows. Protein concentration was quantified using absorbance at 280 nm (NandDrop 2000c, Thermo Scientific). 400 μg of protein was then reduced for 1 hour at 60°C with 10mM tris (2-carboxyethy) phosphine (TCEP) and cysteine-blocked with 8mM iodoacetamide at room temperature in the dark for 30 minutes. Samples were then digested with mass-spectrometry grade trypsin (1:20, Trypsin:Substrate, Promega Gold Trypsin, San Luis Obispo, CA) overnight at 37°C [66]. After digestion, samples were acidified and solvent was removed using a Speed-Vac concentrator. Digested peptides were resuspended in sample buffer (MS-grade H_2_O, acetonitrile and TFA, 20:80:0.4 (v/v)) then enriched for phosphopeptides, and purified using graphite columns according to the manufacturer’s instructions (Pierce TiO_2_ Phosphopeptide Enrichment and Clean-up Kit #88301). Enriched samples were then dried a second time using a Speed-Vac concentrator.

### Shotgun Phosphoproteomics LC-MS/MS

MS data collection for the prepared peptide samples was performed with nanospray UHPLC-MS in the Montana State University Mass Spectrometry Core Facility [67–69]. The dried peptides were resuspended in 50 μL of mass spectrometry grade water, acetonitrile, and formic acid (98:2:0.1v/v/v) with shaking for 15 minutes, and transferred to auto sampler vials. Aliquots of 5 μL each were sampled via Dionex Ultimate 3000 nano UHPLC, with an Acclaim PepMap100 C18 column used for trapping (100 μm x 2cm) and an Acclaim PepMap RSLC C18 (75 μm x 50 cm, C18 2 um 100A) for final peptide separation. The loading pump used 97% H_2_O, 3% acetonitrile, with 0.1% formic acid (v/v). The analytical/elution pump solvent consisted of H_2_O with 0.1% (v/v) formic acid for channel “A” and acetonitrile for channel “B”.

Chromatography was as follows: for 10 minutes, the sample was loaded onto the trapping column at a flow rate of 10 μL/min. From 10 to 11 minutes, the loading pump flow rate was ramped down to 5 μL/min. At 12 minutes, the loading valve was switched to place the trapping column in-line with the analytical column, and from 12 to 17 minutes the loading pump flow rate was ramped from 5 to 40 μL/min. From 106 to 108 minutes the loading pump was ramped back from 40 to 5 μL/min. The analytical/elution pump was maintained at 500 nL/min for the entire run. From 12 to 90 minutes the analytical pump solvents were ramped from 5% to 30% B. From 90 to 105 minutes the analytical pump solvents were ramped from 30% to 95% B. At 110 minutes, the loading valve was switched to divert the trapping column to waste. From 119 to 120 minutes the analytical pump solvents were ramped from 95% to 5% B, and at 120 minutes each run was completed.

The mass spectrometer was a Bruker maXis Impact with CaptiveSpray ESI source: resolution is ~40,000 and accuracy is better than 1 ppm. Spectra were collected in positive mode from 150 to 2200 m/z at a minimum rate of 2 Hz for both precursor and fragment spectra, and with adaptive acquisition time for highly abundant ions (16 Hz for >= 25000 counts to 4 Hz for < 2500 counts).

### Data Processing

We converted the resulting data files with MSConvert (ProteoWizard [70]) to 32-bit .mzML format. These files were then processed with a series of bioinformatics tools. OpenMS TOPPAS was used with published methods [71] to create XTandem workflows to search two established databases (SwissProt and TrEMBL [72]) to evaluate all possible matches. Both fixed (f) and variable (v) modifications were considered in the database search (carbamidomethyl modification of cysteine (f), oxidation of methionine (v), phosphorylation of serine (v), phosphorylation of threonine (v), and phosphorylation of tyrosine (v)), allowing for up to two missed cleavage sites. Precursor tolerance was 50 ppm and MS/MS fragmentation tolerance was 0.05 Da [20]. Instrument type was set to ESI-quad-TOF, and peptide charges up to +3 were permitted. The search database included the reviewed, *Homo sapiens* database (Uniprot/Swissprot) modified to contain both targets and “reversed” decoys for FDR (false discovery rate) corrections for multiple comparisons [73, 74]. Protein-level quantification included all isoforms.

Processed data files were then exported into Excel (Microsoft, Redmond, WA) for further processing. Cluster analysis was performed in MATLAB (Natick, MA) using the clustergram function and complete linkage. Pathway analysis was performed IMPaLA [34] and domain analysis was performed using DAVID [38, 39]. The background protein set for searches was Uniprot Release 2019_02.

### Statistical Analysis and Candidate Selection

To assess the effects of mechanical loading on chondrocyte protein phosphorylation, three randomly assigned groups of cell-seeded agarose hydrogels were established: unloaded control samples (UC), samples undergoing 15 minutes of dynamic compression (DL15), and samples undergoing 30 minutes of dynamic compression (DL30). Statistical analysis was performed using the combined samples for each loading group, from each donor (n=5 samples per group). We defined detected phosphoproteins as those present in the majority of samples (*e.g*. more than 60%). To determine loading induced differences between samples, each dataset was compared against the others to determine (1) unique phosphoproteins to each loading group and (2) overlapped or shared phosphoproteins between loading groups. Four separate comparisons were made: (1) UC vs. DL15, (2) UC vs. DL30, (3) DL15 vs. DL30, and (4) UC vs. DL15 vs. DL30. For phosphoproteins identified in all three groups (*i.e*. UC, DL15, and DL30), statistical comparisons were made using Wilcoxon signed rank tests and Kruskal-Wallis oneway analysis of variance.

## Supporting information

Pathway Enrichment Results

Protein Domain Results

Suppmental Table 5

## Acknowledgements

Funding was provided by NIH P20GM10339405S1, NSF 1342420, NSF 1554708, Montana State University, and the Murdock Charitable Trust.

## Supplemental Informations

*Supplemental Table 1*

Signaling pathways determined from pathway enrichment analysis comparing between unloaded controls (UC) and 15 minutes of dynamic compression (DL15).

*Supplemental Table 2*

Signaling pathways determined from pathway enrichment analysis comparing between unloaded controls (UC) and 30 minutes of dynamic compression (DL30).

*Supplemental Table 3*

Signaling pathways determined from pathway enrichment analysis comparing between 15 and 30 minutes of dynamic compression (DL15 and DL30, respectively).

*Supplemental Table 4*

Signaling pathways determined from pathway enrichment analysis comparing overlapping proteins between all conditions.

*Supplemental Table 5*

Complete proteomics dataset. Labels as follows: raw—detected proteins. refined—detected proteins after FDR filtering. majority—detected proteins from a majority of the samples in each group.

*Pathway Enrichment Results*

Zipped directory of all pathway results based on clusters in Figure 3.

*Protein Domain Results*

Zipped directory of all protein structural domains detected and summarized in Tables 3–4.

